# A Zika virus-responsive sensor-effector system in *Aedes aegypti*

**DOI:** 10.1101/2023.02.06.527261

**Authors:** Sanjay Basu, Christine M. Reitmayer, Sarah Lumley, Barry Atkinson, Mathilde L. Schade-Weskott, Sara Rooney, Will Larner, Eugenia E. Montiel, Rafael Gutiérrez-López, Emily Levitt, Henry M. Munyanduki, Ahmed M. E. Mohamed, Andrew T. Clarke, Sandra Koit, Eva Zusinaite, Rennos Fragkoudis, Andres Merits, Luke Alphey

## Abstract

Zika virus (ZIKV) is a recently re-emerged flavivirus transmitted primarily through the bite of an infected mosquito, *Aedes aegypti* being the main vector. ZIKV infection is associated with a range of adverse effects; infection during pregnancy can lead to foetal abnormalities, including microcephaly. Lacking a licensed vaccine, or specific therapeutics, control of ZIKV transmission focuses on vector control. However, in most transmission settings, current methods are insufficient to successfully control ZIKV, or other similarly-transmitted arboviruses such as dengue and chikungunya viruses. This has stimulated interest in genetics-based methods, either to reduce the number of mosquitoes (“population suppression”), or to make mosquitoes less able to transmit (“population modification”). Here, we describe a method to selectively eliminate infected mosquitoes, using a virus sensor inserted into the mosquito genome and coupled to a quorum-counting lethal effector. In mosquitoes, ZIKV normally establishes persistent, lifelong infection; survival of these infected mosquitoes is crucial to transmission potential. Correspondingly, removal of infected mosquitoes can reduce vectorial capacity of a mosquito population, i.e. ability to transmit. Since relatively few mosquitoes become infected, typically <2%, engineered hypersensitivity to ZIKV would have only a modest population-level fitness cost, and lower still if transmission were successfully reduced by such means.

## Main text

Population modification approaches have focused on making mosquitoes less permissive to pathogen replication, such that little or no pathogen reaches the salivary glands and saliva, thereby reducing transmission. However, we reasoned that hypersensitivity, such that infected mosquitoes become incapacitated and unable to transmit the virus, would be equally effective in preventing onwards transmission. Ideally, the effect would be rapid, so that the mosquito is incapacitated before the mosquito becomes infectious. The time from ingesting an infected blood meal to becoming infectious (Extrinsic Incubation Period, EIP) varies with temperature and between different viruses; for Zika virus (ZIKV) in *Ae. aegypti* the EIP is around 7 days (6-9 days at 28°C)^1^. This is a significant fraction of the adult female lifespan; life-shortening has previously been proposed as a powerful method to reduce transmission^2, 3^.

In concept, a synthetic genetic circuit providing a response to infection might comprise a sensor for infection, coupled to an effector that is expressed, or de-repressed, when the cell is infected, thereby activating the sensor. Though cells and mosquitoes mount their own responses to infection, implying natural, in-built sensor systems, these do not appear sufficiently specific for our purpose. We therefore set out to build a synthetic sensor. Infected cells contain several virus-encoded molecules and enzymatic activities that are absent from uninfected cells. We focused on the NS2B/NS3 protease, which is required to process the virus polyprotein at well-defined sites not cleaved by cellular proteases. Previous studies have shown that recombinant proteins engineered to contain suitable cleavage sites can be selectively cleaved by virus-encoded protease, using proteases and corresponding cleavage sites from several viruses including DENV, leading to modest de-repression or translocation of a fluorescent reporter in transfected cells^4-8^.

To provide a flexible means of detecting virus-induced proteolytic cleavage, we constructed “tethered-Cre” sensors (Fig 1a, b and d). Cre recombinase, a tyrosine recombinase from the P1 bacteriophage, catalyses site-specific recombination between two DNA recognition sites (*lox*P sites). We fused Cre to a cytoplasmic tether via a cleavable linker (Fig 1a and c). By excluding it from the nucleus, this renders Cre functionally inert until the linker is cleaved, at which point Cre can translocate to the nucleus and interact with DNA targets. A design following this principle in mammalian cells demonstrated strong induction of an eGFP reporter following DENV infection^4^. However, 2-3% of uninfected cells also expressed eGFP, which might be problematic in mosquitoes if using a lethal effector in place of a neutral reporter. We therefore used a novel “double-tether” design using a transgene expressing a section of the ZIKV non-structural proteins with Cre recombinase encoded in place of NS3, between NS2B and NS4A, resulting in Cre being tethered to the endoplasmic reticulum (ER) membrane by fusion to transmembrane domains at both N- and C-termini. The NS2B/NS3 protease cleavage sites (“linkers”) naturally present at the NS2B-NS3 and NS3-NS4A junctions were preserved. ZIKV polyprotein is the key natural substrate of viral protease so these linkers are predicted to be presented in a relevant subcellular and molecular context. Cre is released only when both linkers are cut by the NS2B/NS3 protease, present only in infected cells.

**Fig 1:**
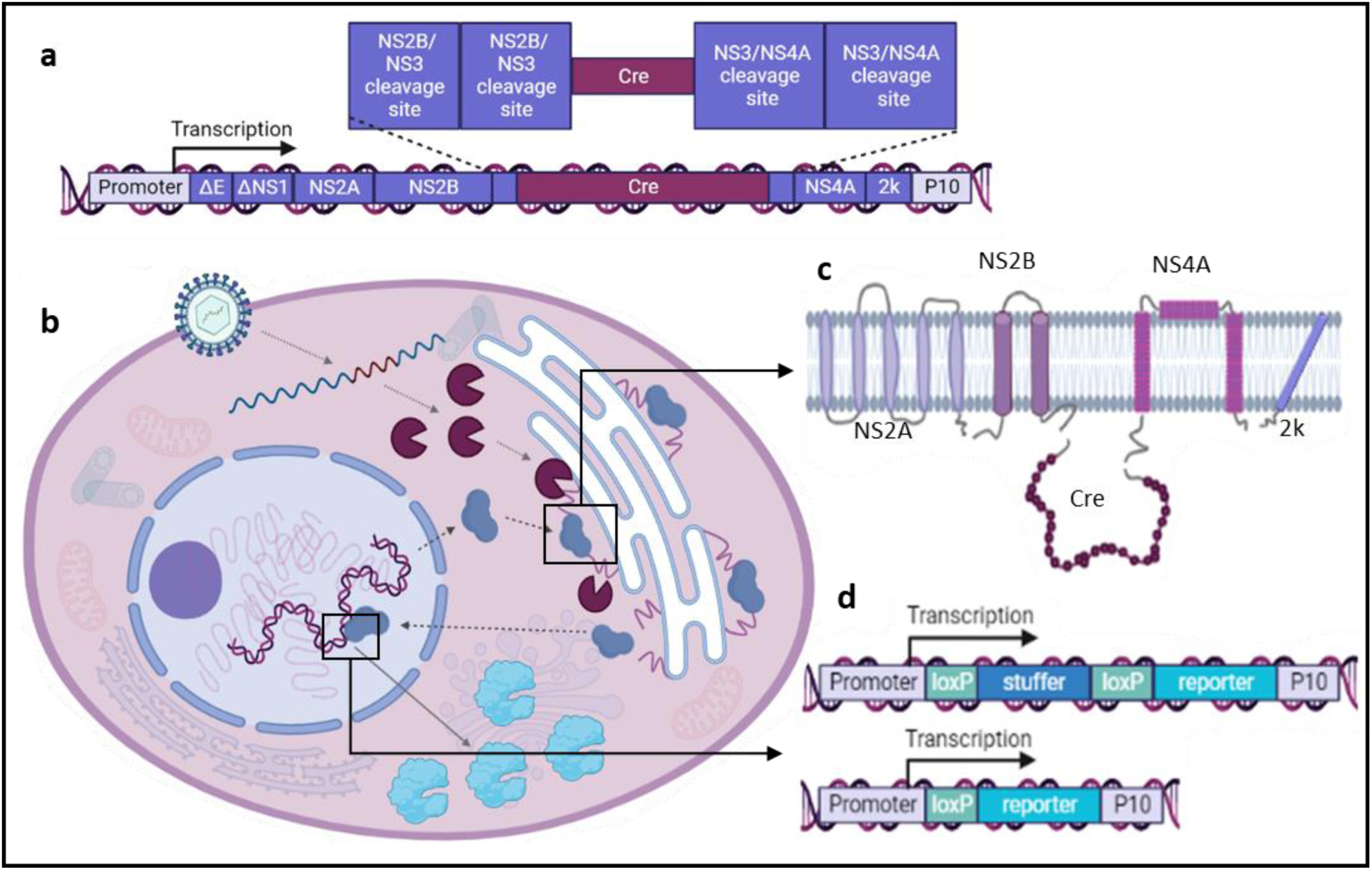
**a. Tethered-Cre construct (“ZZ-Cre”).** Polyubiquitin promoter drives expression of a derivative of the ZIKV polyprotein comprising parts of the E and NS1, then NS2A through to 2k. Cre replaces the NS3 sequence and is flanked by two repeats of NS2/NS3 protease cleavage site at the N-terminal end and two repeats of NS3-NS4A protease cleavage site at the C-terminal end. **b. ZIKV infected cell expressing ZZ-Cre and loxP reporter construct**. Modified cells express ZZ-Cre, which is tethered to the endoplasmic reticulum (ER, dashed arrow) by fusion to transmembrane domains derived from ZIKV polyprotein (**c**). When the cell becomes infected by ZIKV NS2B/NS3 protease is expressed. This cleaves protease-sensitive sequences flanking Cre, releasing it from the ER tether. Cre translocates to the nucleus (dashed arrows). In the nucleus, Cre cause recombination of the loxP reporter/effector construct, causing a permanent switch from expression of the stuffer fragment protein to the reporter/effector protein (**d**, solid arrow).

If released from its tether by NS2B/NS3 protease, Cre is free to translocate to the nucleus, where it can catalyse recombination between engineered *lox*P sites of an effector transgene inserted into the mosquito genome. For initial experiments in mosquito cell culture, we constructed a Cre-inducible luciferase reporter. This comprises a “stuffer fragment”, encoding mCherry and flanked by *lox*P sites, between a strong, ubiquitous promoter (polyubiquitin)^9^ and a sequence encoding nanoluciferase (NLuc, Fig 1d). Additional stop codons were included between NLuc and *lox*P to minimise basal expression of the reporter prior to activation by *lox*P recombination. In the absence of NS2B/NS3 protease, NLuc expression in cells with tethered-Cre sensors and Cre-responsive *lox*P-NLuc reporter was similar to non-transfected cells, i.e. close to assay/instrument background, but were >1000-fold higher in the presence of a ZIKV replicase-encoding plasmid, which expresses all ZIKV non-structural proteins, including NS2B-NS3 protease (Fig 2a). This signal was about 16-fold lower than from a positive control plasmid comprising the NLuc reporter with the stuffer fragment removed, indicating that ∼6% of *lox*P-reporter plasmids were activated by Cre in these experiments. We compared a construct with tandem duplication of the cleavage sites (ZZ-Cre) with the native configuration (Z-Cre) of one cleavage site each side of Cre. For both Z-Cre and ZZ-Cre we observed a significant induction of NLuc expression in the presence of a ZIKV replicase (Z-Cre: p = <0.0001, t (34) = 4.723; ZZ-Cre: p 0.0025, t (34) = 3.272). There was no significant difference in induced expression levels between the Z-Cre and ZZ-Cre design in the absence of replicase, however, background expression levels in the Z-Cre design were slightly higher compared to baseline levels of the ZZ-Cre design (p = 0.0103, t (34) = 2.717). Based on these results, we used the ZZ-Cre design for mosquito infection studies. In cell culture we also tested ZZ-Cre/*lox*P-NLuc for responsiveness to NS2B/NS3 proteases originating from a range of other flaviviruses and found no significant induction of NLuc expression in the presence of the replicase of insect-specific Niénokoué virus (NIEV) but significant expression induction with replicase from dengue virus (DENV, p = 0.0247, t (10) = 2.641) and Kunjin virus (KUNV, p = 0.0128, t (10) = 3.025). This indicates that broad spectrum application of this sensor might be a feasible goal for the future, also that cross reaction with non-pathogenic insect-specific flaviviruses could potentially be avoided (Fig 2b).

**Fig 2:**
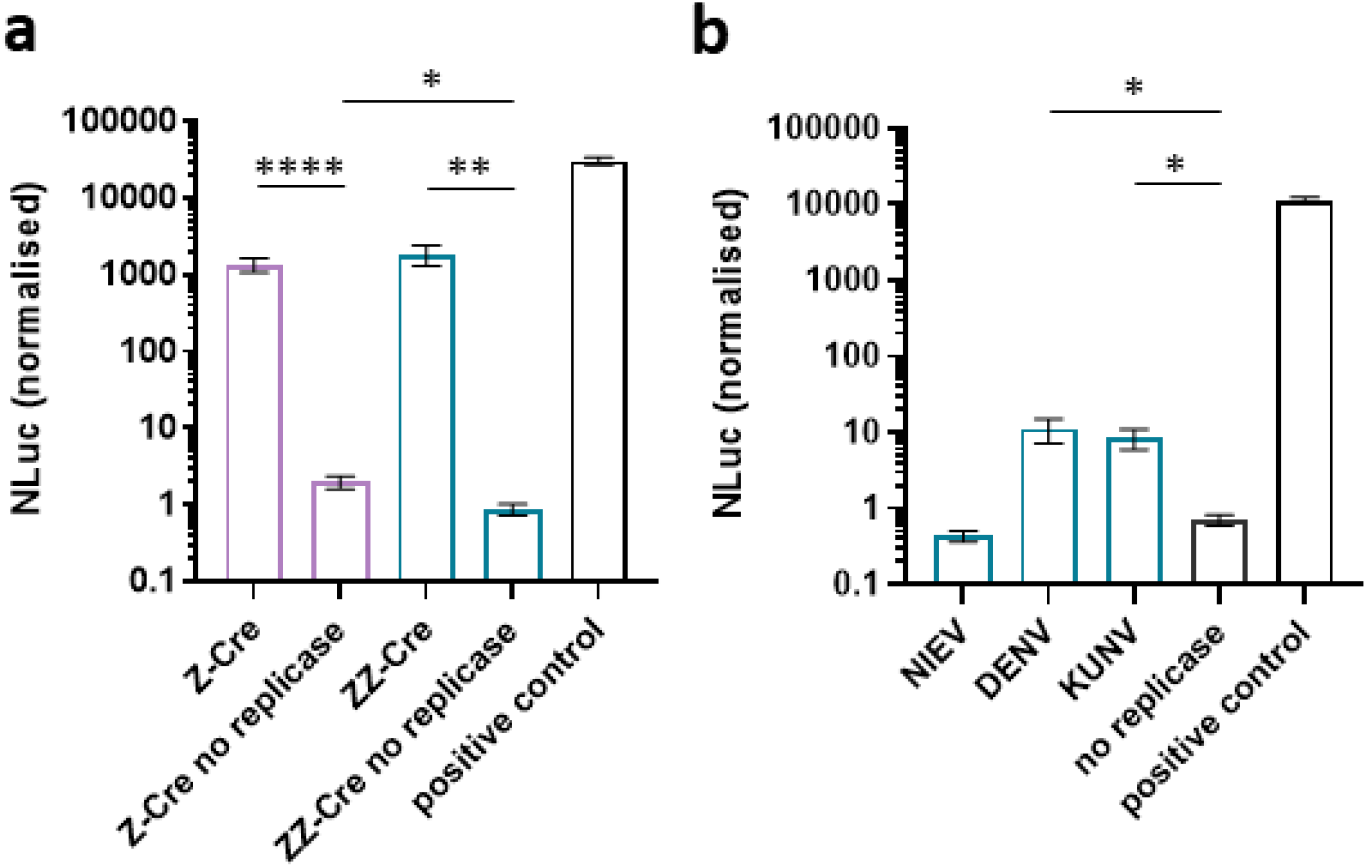
**a. Tethered-Cre designs induce strong expression of NLuc in presence of ZIKV replicase.** NLuc expression levels (normalized against a firefly luciferase transfection control) of the Z-Cre and ZZ-Cre tethered-Cre design against ZIKV replicase. Both Z-Cre and ZZ-Cre tethered-Cre designs provide increased NLuc expression in the presence of ZIKV replicase (source of NS2B-NS3 protease). A fully recombined NLuc expression plasmid was used as positive control to indicate the maximum achievable response. Data shown are NLuc expression normalised against a firefly luciferase control. **b. Response of ZZ-Cre tethered-Cre design to NIEV, DENV and KUNV replicases**. In the presence of NIEV replicase, no induction of NLuc expression was observed; presence of both DENV and KUNV replicase resulted in NLuc expression induction. Significant differences are indicated with asterisks (*: p<0.05, **: p<0.01, ****: p<0.0001).

Translating a sensor-effector system to live mosquitoes is dependent on many factors including the proportion of cells that become infected with ZIKV upon infection, the effector response required to incapacitate the mosquito, and fitness costs from uninduced activation. We therefore developed a novel “quorum-counting” mechanism based on expression of a secreted toxin such that very low expression, e.g. from a very small number of cells, should be harmless but expression above a threshold level should be lethal. We used AaHIT (*Androctonus australis hector* insect toxin), an arthropod-specific peptide toxin originally from a scorpion venom. AaHIT affects neuro-muscular junctions; in a previous study, transient expression of AaHIT in *Ae. aegypti* induced transient knock-down^10^. Here, *lox*P recombination and removal of the stuffer fragment in infected cells leads to permanent induction of AaHIT. Importantly, AaHIT does not kill the expressing cells, so these, and overall AaHIT expression, can increase as infection spreads until the threshold for a physiological effect is reached.

The limited prior use of Cre-*lox*P in *Ae. aegypti* focused on germline recombination, which was readily observed but not especially efficient (e.g. 33% of families showed at least one recombination event following injection of a Cre-expressing plasmid)^11^. We therefore investigated the use of Cre-*lox*P in somatic cells of transgenic mosquitoes. We replaced NLuc with AmCyan, to generate AmCyan fluorescence after Cre-induced recombination of *lox*P and inserted this PUb-*lox*P-mCherry-*lox*P-AmCyan (“*lox*P-AmCyan”) transgene into *Ae. aegypti*. The PUb promoter used is expected to give strong expression throughout the mosquito and was used for both sensor and effector components^9^. We exposed the resulting *lox*P-AmCyan mosquitoes to Cre, by crossing to mosquitoes expressing Cre (tPUb-Cre) or injecting tPUb-Cre plasmid (Fig 3). In the presence of Cre, blue fluorescence was readily observed, indicating Cre-dependent recombination and expression of the AmCyan ‘effector’.

**Figure 3:**
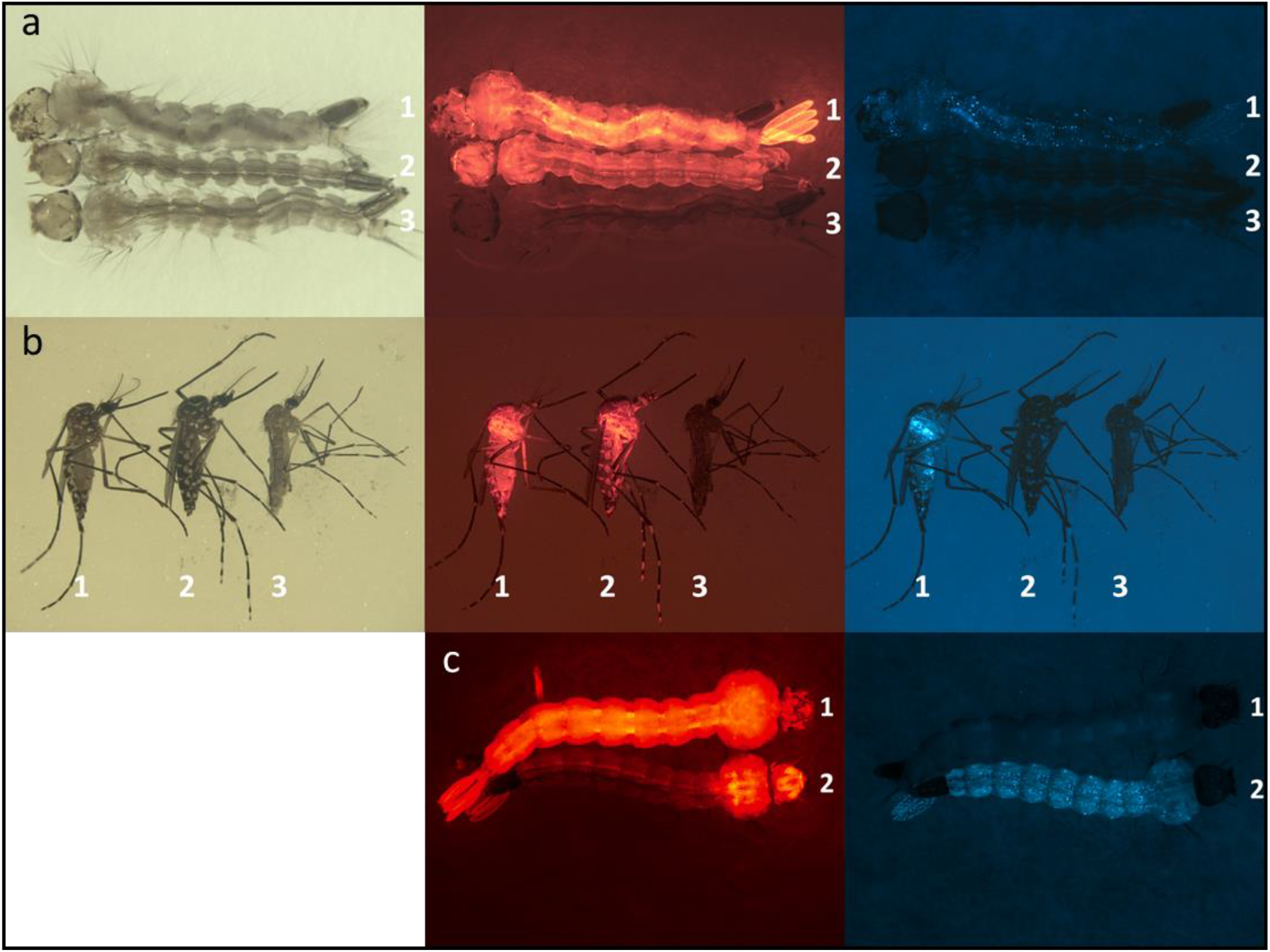
Cre stimulates recombination and AmCyan expression in *lox*P-AmCyan transgenic mosquitoes. *lox*P-AmCyan transgenic mosquitoes were crossed to transgenic mosquitoes carrying tPUb-Cre. tPUb is a truncated version of the Pub promoter giving weak, ubiquitous expression. Progeny carrying both constructs were analysed by fluorescence microscopy. *lox*P-AmCyan expresses mCherry unless subject to *lox*P recombination, in which case it expresses AmCyan, providing red or cyan fluorescence, respectively. Larvae (a) and adults (b) show cyan fluorescence only in the presence of both transgenes (1: both transgenes, 2: *lox*P-AmCyan only, 3: tPub-Cre only). c: *lox*P-AmCyan embryos were injected with tPUb plasmid at the posterior pole. Survivors were observed as larvae. Relative to non-injected control (1: no plasmid, 2: tPUb Cre plasmid), injected larvae showed expression of AmCyan in abdomen and thorax, and loss of mCherry expression in posterior regions.

We next generated PUb-*lox*P-mCherry-*lox*P-AaHIT (“*lox*P-AaHIT”) transgenic mosquitoes to evaluate the same system with a lethal effector. This construct was intended to provide AaHIT expression only after Cre-induced removal of the stuffer fragment. Simple breeding experiments revealed no obvious fitness defects, e.g. from unintended basal expression of AaHIT (data not shown). To test the function of the *lox*P-AaHIT transgene we crossed *lox*P-AaHIT transgenic mosquitoes to tPUb-Cre mosquitoes. No offspring carrying both *lox*P-AaHIT and tPUb-Cre were recovered (Table S2), indicating that Cre-induced mosaic expression of AaHIT was lethal, as anticipated (Figure S1). However, death occurred early in development, whereas for bloodmeal-related transmission of ZIKV, the relevant sex and developmental stage is adult females. We therefore investigated the phenotype of *lox*P-AaHIT when exposed to Cre in adult females. We used a Cre-expressing Semliki Forest virus, SFV-Cre, to express Cre in infected cells of adult females. This approach simulates an ideal tethered-Cre sensor, in which all virus-infected cells produce untethered Cre, but other cells do not. Alphaviruses such as SFV can be engineered to express exogenous proteins from their subgenomic promoter^12^, though SFV-Cre was found to be unstable with limited Cre expression after 2 passages. Infection of AaHIT mosquitoes by intrathoracic inoculation with 5×10^4^ and 5×10^3^ PFU/mosquito of SFV-Cre induced >90% mortality by day 5 post injection, while at 5×10^2^ PFU/mosquito around 50% survived (Fig 4a). Comparing survival of individuals injected with SFV-Cre or similar amounts of SFV-GLuc – expressing a *Gaussia* luciferase reporter instead of Cre^13^ – we found a significant difference (χ2=187.4, P<0.0001) between these groups. Within each treatment dose, we found a significant difference in survival between individuals injected with SFV-Cre and individuals injected with SFV-GLuc (for 10^4^: χ2=61.76, P<0.0001; for 10^3^: χ2=38.91, P<0.0001; for 10^2^: χ2=17.59, P<0.0001) with female mosquitoes injected with SFV-Cre having significantly reduced survival compared to SFV-GLuc injected females. Within the SFV-Cre injected group, there was a significant dose dependent difference in survival (χ2=70.58, P<0.0001); there was no dose dependent difference in survival within the SFV-GLuc group (χ2=5.070, P=0.0793). We compared recombination rates in gDNA extracts of individuals by PCR from the SFV-Cre treatment group to the SFV-GLuc treatment group and found a significantly higher recombination rate in the SFV-Cre treatment group (Fig 4b, Mann–Whitney U = 4, n1 = 13, n2 = 4, P = 0.0101, two-tailed).

**Fig. 4:**
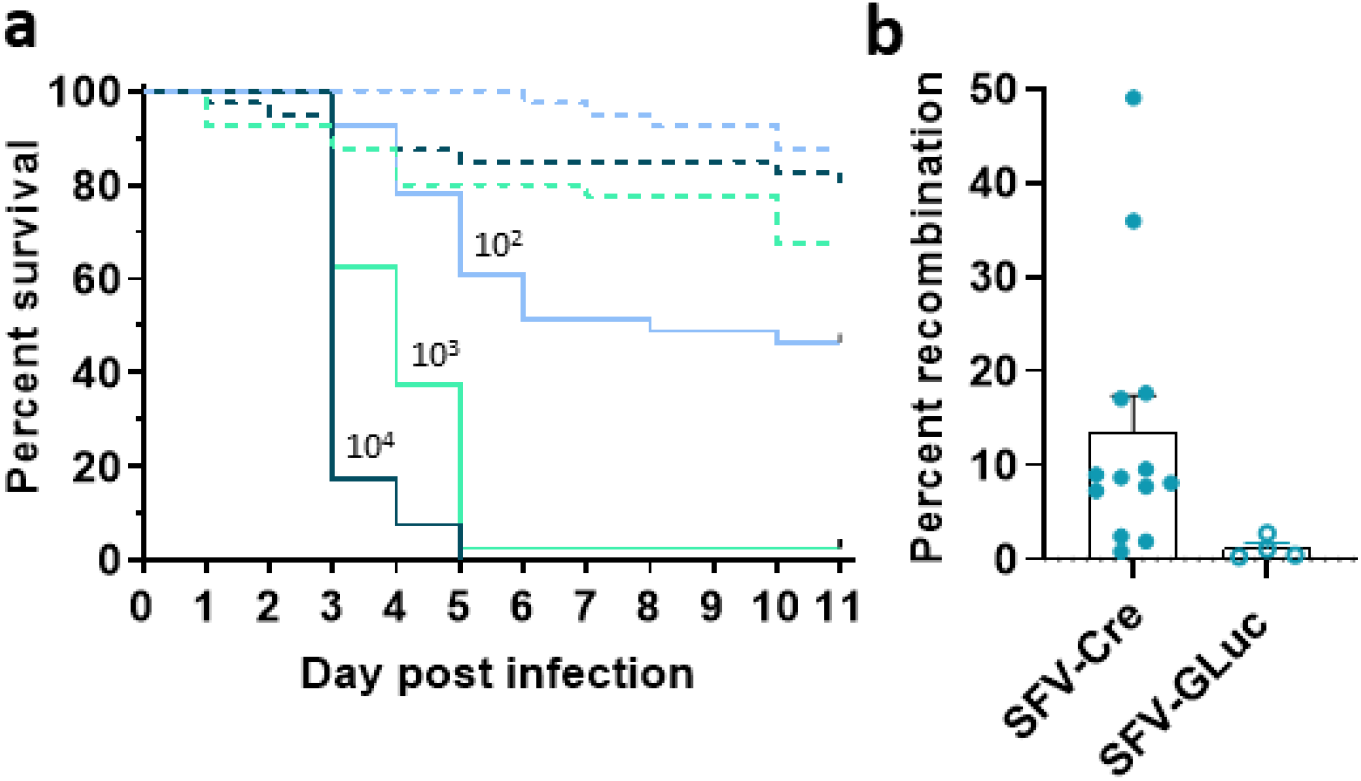
**a. Survival of female mosquitoes injected with SFV-Cre or SFV-GLuc at three different doses (10^4^, 10^3^ or 10^2^ PFU per mosquito).** Survival was significantly lower for SFV-Cre injected mosquitoes (solid lines) compared to SFV-GLuc injected mosquitoes (dashed lines, for 10^4^: χ^2^=61.76, P<0.0001, for 10^3^: χ^2^=38.91, P<0.0001, for 10^2^: χ^2^=17.59, P<0.0001, n(ZZ-Cre/AaHIT ZIKV) = 128, n(ZZ-Cre/AaHIT DMEM) = 80, n (LVP ZIKV) = 80, n (LVP DMEM) = 80. There was a dose dependent reduction in survival within the SFV-Cre group with treatment groups being injected with higher titres having a worse survival outcome compared to groups with lower injection titres (10^4^ PFU/mosquito in dark green, 10^3^ PFU/mosquito in light green, 10^2^ PFU/mosquito in light blue). Within each dose treatment, groups that had been injected with SFV-Cre showed a significantly reduced survival compared to groups injected with SFV-GLuc. **b. loxP recombination rate in gDNA of mosquitoes infected with SFV-Cre and SFV-GLuc**. Mosquitoes infected with SFV-Cre were analysed to estimate the ratio of recombined to non-recombined *lox*P-AaHIT transgenes; loxP recombination was observed up to 49% (mean 13.5%) in SFV-Cre injected mosquitoes. In comparison to SFV-Gluc injected mosquitoes (mean 1.1% recombination) this is a significantly higher rate of *lox*P recombination as measured by qPCR (Mann– Whitney U = 4, n1 = 13, n2 = 4, p = 0.0101, two-tailed, shown as individual datapoints with mean and SEM).

Next we combined the two components, tethered-Cre sensor and *lox*P-AaHIT lethal effector, to provide a prototype complete sensor-effector system. We generated transgenic mosquitoes carrying a transgene expressing tethered ZZ-Cre. Tethered ZZ-Cre and *lox*P-AaHIT insertion lines were crossed and then bred to homozygosity for both insertions - this provides a true-breeding line (“ZZ-Cre/AaHIT”). We reasoned that higher copy number of sensor and effector would likely improve sensitivity. It could also have increased fitness costs through increased expression of system components, elevated uninduced recombination of *lox*P-AaHIT, or insertion mutation effects, which are typically recessive. The ZZ-Cre/AaHIT mosquitoes were viable and fertile, but more detailed studies would be required to identify quantitative fitness effects, if any.

We then infected both ZZ-Cre/AaHIT mosquitoes and wildtype (Liverpool strain, LVP) mosquitoes with ZIKV by intrathoracic (IT) inoculation Fig 5, solid lines) together with injecting a different subset of mosquitoes of these same groups with DMEM (Fig 5, dashed lines) as a no-infection negative control. IT inoculation provides ∼100% infection rates and consistent titres in the infected mosquitoes^14^. We found a statistically significant difference in survival between the four different treatment groups (χ2=524.3, P <0.0001). Pairwise comparison revealed that the difference in survival is highly significant between ZZ-Cre/AaHIT individuals injected with ZIKV compared to those injected with DMEM (χ2=191.9, P <0.0001). There was no difference in survival between uninfected (DMEM-injected) ZZ-Cre/AaHIT and LVP individuals (χ2=1.699, P=0.1925). Inoculation with ZIKV had no effect on survival in LVP individuals (χ2=2.236, P=0.1348). While all control groups showed >80% survival over the duration of the experiment (8 days post injection), ZIKV infected ZZ-Cre/AaHIT mosquitoes had a median survival of only 5 days post-inoculation, with none surviving beyond 7 days (Fig 5). This indicates a fully-penetrant lethal effect from the sensor-effector system following infection with ZIKV.

**Fig. 5:**
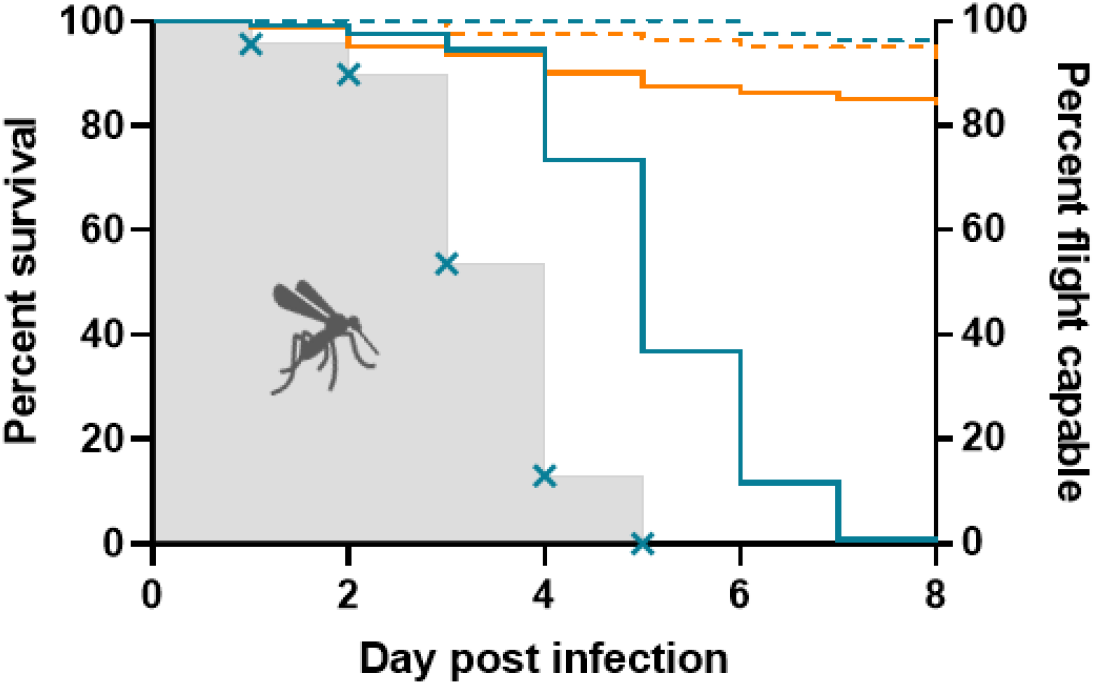
Survival of ZZ-Cre/AaHIT transgenic mosquitoes after IT infection challenge with ZIKV. Transgenic mosquitoes (blue line) infected with ZIKV (solid line) showed significantly reduced survival compared to uninfected (PBS injected, dashed line) ZZ-Cre/AaHIT transgenic mosquitoes as well as compared to infected and uninfected LVP mosquitoes (orange, solid and dashed lines, respectively). The proportion of transgenic, ZIKV infected mosquitoes showing normal movement is indicated by blue crosses; the grey area indicates the proportion of flight-capable females over time.

During these experiments, we observed ZIKV infected ZZ-Cre/AaHIT mosquitoes showing uncoordinated and semi-paralysed behaviours prior to death. This distinctive syndrome resembled effects we previously observed from transient expression of AaHIT^10^. Death of an infected mosquito clearly prevents onward transmission of virus via biting, but death is not strictly required; anything that prevents biting would have a similar effect. Inability to fly properly would prevent the mosquito’s normal host seeking, blood feeding, and predator evasion behaviours. We therefore additionally scored gross flight defects: inability to fly the 10 cm to the top of their container. By day 5, none of the ZIKV infected ZZ-Cre/AaHIT mosquitoes still alive were flight capable (Fig 5, grey shaded area, blue crosses, show flight capable mosquitoes as proportion of infected mosquitoes).

In these experiments, our sensor-effector system provided 100% mortality after seven days following infection with ZIKV. No obvious deleterious effects were observed in the absence of ZIKV infection, though additional studies would be required to identify more subtle fitness effects, if any. For further use, a panel of insertion lines carrying the complete system could be generated, from which ones with high effector penetrance and low fitness costs could be selected. Fitness costs are likely important for any future field use, since significant fitness cost would tend to lead to the system reducing in allele frequency over time. In this context, the key consideration is basal fitness cost, which applies to all carriers. The fitness cost induced by infection is, perhaps surprisingly, less significant – few wild mosquitoes become infected with ZIKV and other flaviviruses. Similar to dengue virus, mosquito surveillance typically finds a relatively low proportion of adult female mosquito vectors positive for ZIKV RNA even in areas of active transmission (e.g. 0-9% in various settings)^15-19^. Since these data may not distinguish infected mosquitoes from those merely containing virus RNA, e.g. in their midgut from a recent blood meal, the proportion of infected mosquitoes is presumably lower still. Furthermore, modifying a significant proportion of a vector population to this hypersensitive phenotype should greatly reduce the vectorial capacity of the population – and correspondingly reduce transmission and hence the proportion of mosquitoes becoming infected and incurring that fitness cost. More speculatively, such a system may introduce a selective advantage to mosquitoes that avoid biting virus infected hosts, though it is not clear whether this trait is amenable to selection.

Various protease-based sensors have been demonstrated in mammalian tissue culture cells^4-8^. These imply that our design could readily be adapted to other important mosquito-borne flaviviruses including dengue virus and yellow fever virus, simply by changing the protease-sensitive linker sequence. It is further likely that these linkers can be inserted as tandem repeats, as we demonstrated in duplicating the ZIKV cleavage sites, thereby giving a single sensor that can respond to any of a predetermined set of viruses.

While such sensor-effector systems could be introgressed into wild vector populations simply by releasing large numbers to interbreed (“inundative release”), this is unlikely to be cost-effective for large-scale interventions, particularly if the transgene does have significant fitness costs. The obvious alternative is to introgress the system with a “gene drive” system. Gene drives can increase in frequency and persist in the target population despite some fitness cost to individual carriers^20-22^. No engineered gene drives have yet entered field use, but various prototypes have been successfully demonstrated in laboratory cages. Engineered hypersensitivity systems such as we have demonstrated might make ideal “cargo” elements for such gene drives.

## Methods

### Plasmid construction

Cre, firefly luciferase (FLuc) and loxP reporter/effector plasmids were based on high-copy-number ColE1 origin plasmid as a backbone. Used promoters, terminators and encoded proteins are shown in Figure S1. Sequences are available on Genbank.

Virus replicase plasmids: Replicase plasmids or ZIKV, KUNV, DENV and NIEV were based on the pCCI-Bac1 inducible-copy-number plasmid as a backbone. The replicases were expressed using the insect-specific polyubiquitin promoter; the coding sequences included region encoding for the 35 C-terminal amino acids of E protein (to ensure correct anchoring of NS1 protein to the membranes), following by the full-length non-structural polyprotein NS1-NS5 of respective viruses. The open reading frame terminated with the stop codon and SV40 poly(A) signal was used as a transcriptional terminator.

### Cell lines

Vero cells (ECACC 84113001) were maintained in Dulbecco’s Modified Eagle Medium (DMEM, Life Technologies), supplemented with 10% heat-inactivated fetal bovine serum (FBS, Gibco) and 10% tryptose phosphate broth (TPB, Gibco) at 37°C and 5% CO_2_. BHK-21 baby hamster kidney cells (CLL-10, ATCC) were cultured in Glasgow’s minimal essential medium (Life Technologies), supplemented with 10% heat-inactivated FBS and 10% TPB and 100 U/ml penicillin (Gibco) and 0.1 mg/ml streptomycin (Gibco) at 37°C and 5% CO_2_. *Aedes aegypti* derived Aag2 cells were maintained in Leibovitz’s L-15 medium (Life Technologies), supplemented with 10% heat-inactivated FBS, 10% TPB, 100 U/ml penicillin and 0.1 mg/ml streptomycin at 28°C.

### Tethered-Cre/loxP NLuc replicase transfection experiment

Aag2 cells were seeded into 96-well plates at a density of 40,000 cells/well. The next day, cells were transfected with a DNA complex mixture consisting of *lox*P-NLuc plasmid (1 ng/well), either tethered Z-Cre or ZZ-Cre expression plasmid (1 ng/well) and the relevant ZIKV, NIEV, DENV or KUNV replicase expression plasmid (10 ng/well), as well as relevant controls such as no replicase negative control and a fully recombined loxP NLuc plasmid (1 ng/well). All treatment conditions included an internal FLuc expression plasmid (0.2 ng/well) as normalization control. Transfection was performed using the TransIT-PRO® Transfection Reagent & Kit (Mirus Bio). Cells were incubated at 28°C following transfection for 48h. Afterwards, cells were washed twice in PBS, lysed in passive lysis buffer (Promega) and luciferase assays were performed using the Nano-Glo® Dual Reporter Assay Kit (Promega) and the GloMax®-Multi Detection luminometer System (Promega). The experiments were carried out in three biological replicates for the Z-Cre vs ZZ-Cre ZIKV replicase comparison experiment and once for the ZZ-Cre NIEV, DENV, KUNV replicase experiment. All biological replicates were carried out in 6 technical replicates per treatment.

### Mosquito transgenesis

Transgenic mosquito lines were generated as previously described^10, 23, 24^. In short, *Ae. aegypti* pre-blastoderm embryos were microinjected with a solution comprising 300 ng/μl of plasmid expressing hyperactive piggyBac transposase ^25^ under control of *Aedes* PUb promoter and 500 ng/μl of donor piggyBac transgene plasmid (as appropriate). After hatching the injected eggs, injection survivors were backcrossed to wildtype (Liverpool strain, LVP) individuals and progeny screened for the fluorescent transformation marker using a fluorescent stereomicroscope (Leica). Single male transformants (G_1_) were then crossed to five LVP females each to establish independent transgenic lines. Insertion-site identification was performed as previously described^26^. To generate homozygotes for both AGG1894 and AGG1595, single adult crosses were setup, and screened for homozygosity using insertion-site specific PCR using non-lethal sampling of one leg per mosquito. Homozygous progeny were self-crossed and each generation was screened for their fluorescence profile.

Transgenic and LVP *Ae. aegypti* were reared and maintained at 26.5°C (±1°C) and 65% (±10%) relative humidity with a 12:12 hour light:dark cycle. Larvae were fed on finely ground TetraMin Ornamental Fish Flakes (Tetra GmbH), and adults were provided with 10% sucrose solution. For generation of egg stocks, adult females were fed on defibrinated horse blood (TCS Biosciences).

### Virus stocks

ZIKV MP1751 stocks were generated by inoculating Vero cells at 80% confluency with ZIKV MP1751 (National Collection of Pathogenic Viruses catalogue number 1308258v) at a MOI of 0.01 and culturing cells for 48h at 37°C and 5% CO_2_. SFV-Gluc and SFV-Cre were rescued from icDNA clones. SFV-Cre had a similar design to the previously published SFV-Gluc virus^13^ except it contained the sequence of Cre under a duplicated viral subgenomic promoter placed downstream of the region encoding for structural proteins. The icDNA plasmid, containing a CMV promoter, was electroporated (pulsed twice using a square wave of 850 volts for 25 milliseconds) into BHK-21 cells using a Gene Pulser Xcell electroporator (BioRad); transfected cells were cultured in a T175 for 48h at 37 °C. The resulting culture media was clarified using centrifugation. ZIKV stocks were also column concentrated using Amicon™ 100 kDa Ultra-15 Centrifugal Filter Units (Merck).

### Mosquito infection through IT inoculation

Viral infection was parenteral in method and followed a standard protocol^14^. Briefly, mosquitoes of 5-7 days-post eclosion were knocked down with CO_2_ and then transferred to a pre-chilled glass petri dish on ice. A prepared glass capillary needle (Drummond Ltd) was filled with the respective inoculum. For SFV-Cre and SFV-Gluc injections mosquitoes received either 5×10^4^, 5×10^3^ or 5×10^2^ PFU/mosquito. For ZIKV infections inoculum at 2.25×10^8^ PFU/ml was used, with each mosquito receiving ∼0.5μl and therefore ∼1.13×10^5^ PFU of virus. Using a dissecting stereomicroscope, the needle impaled a mosquito that was then placed within an adapted carton for observation and data collection throughout the rest of the experiment. Mosquito cartons were placed within a secure environmental incubator (Percival) with conditions of: 28°C (+/-1°C), 80% RH and 12:12 light:dark cycle. The mosquitoes were sustained using cotton wool wads dampened with 10% sucrose solution. Inoculated mosquitoes were observed daily, and any dead (non-mobile) were removed. Mosquitoes were classified incapacitated (non-flying) if they appeared to be motionless, but after tactile stimulation responded with minor movement such as movement of legs (when lying on their back), uncoordinated whole-body movements or small jumps. No mosquitoes classified as incapacitated were able to fly or crawl up the 10 cm walls of their containers.

### qPCR for determination of recombination rate in gDNA

Genomic DNA extraction was performed by homogenizing individual mosquitoes in 180 µl ATL buffer (Qiagen). Following homogenization, 20 µl of proteinase K (Qiagen) was added to each homogenate followed by incubation at 56°C for 2 hours. Genomic DNA was then further purified from the homogenates using the DNeasy Blood and Tissue kit (Qiagen) as per the manufacturer’s instructions, except that samples were eluted in 100 µl of nuclease-free water.

All qPCR reactions were set up using 7.5 µl of 2 x Luna Universal qPCR master mix (NEB), 0.375 pmol forward primer, 0.375 pmol reverse primer, 25 ng of genomic DNA and nuclease free water up to a final volume of 15 µl. For each sample, three targets were amplified, each in triplicate. The first two targets were the non-recombinant and the recombinant versions of loxP-AaHIT as well as GAPDH which was used as a reference gene to normalise the reactions. The two loxP-AaHIT targets used the same forward primer (loxP-AaHIT fwd) in combination with the appropriate reverse primer (either loxP-AaHIT non-recom rev or loxP-AaHIT recom rev) according to the intended target (Table S1). All amplifications were performed under the same thermal cycling conditions using a QuantStudio 3 real-time PCR system (Applied Biosystems). Initial denaturation was performed at 95°C for 1 min, followed by 40 cycles of denaturation at 95°C for 15 s and priming and extension at 60°C for 15 s at the end of which a fluorescence was measured. This was followed by a melting curve from 60°C up to 95°C. Samples were run alongside standards of know copy numbers of non-recombinant and recombinant loxP-AaHIT template. QuantStudio 3 Design and Analysis software (v1.5.1; Applied Biosystems) was used to determine the Ct values for each of the reactions as well as extrapolate the copy number of non-recombinant and recombinant loxP-AaHIT in each mosquito.

### Statistical analysis

Analysis of Z-Cre and ZZ-Cre transfection experiments was carried out using Student’s two-tailed test to compare treatment groups of interest. Analysis of the SFV-Cre survival study was carried out using a Log-rank (Mantel-Cox) test. Recombination qPCR data was analysed using a non-parametric, two tailed Mann–Whitney U test. ZIKV infection survival experiment analysis was carried out using a log-rank test (Mantel-Cox) for overall comparison and Chi square tests for pairwise comparisons. All analysis was carried out using GraphPad Prism 9.3.1.

## Acknowledgements

Figures 1 and S1 were created with BioRender.com. This research was developed with funding from the Defense Advanced Research Projects Agency (DARPA). The views, opinions and/or findings expressed are those of the author and should not be interpreted as representing the official views or policies of the Department of Defense or the U.S. Government.

## Author contributions

Methodology: SB, CMR, SL, BA, RF, MLSW, AM, LA; Validation: SB, CMR, SL, BA, RF, MLSW; formal analysis: CMR; investigation: SB, CMR, SL, BA, MLSW; sample preparation and data collection: SB, CMR, SL, BA, MLSW, SR, WL, EEM, EL, RGL, HMM, AMEM, ATC; writing (original draft preparation): CMR and LA; writing (review and editing): SB, CMR, SL, BA, RF, MLSW, AM, LA; visualization: CMR; project supervision: SB, LA, CR and AM; resources: SK, EZ and AM; funding acquisition: AM, RF and LA. All authors have read and approved the submitted version of the manuscript.

The authors declare no conflict of interest. The funders had no role in the design, execution, interpretation or writing of the study.

## Materials & Correspondence

Material requests and correspondence should be addressed to the corresponding author, Luke Alphey.

## Notes

### Competing Interest Statement

The authors have declared no competing interest.

