## Supplementary information for "A Zika virus-responsive sensor-effector system in *Aedes aegypti*"

**Table S1**: Sequences of primers used in qPCR analysis

| Primer | Sequence (5'→3') |
|---|---|
| *lox*P-AaHIT fwd | TCAAGCGTTTCCTCGTTTCT |
| *lox*P-AaHIT non-recombined rev | TCCTTGATGATGGCCATGTTAT |
| *lox*P-AaHIT recombined rev | GGTGTGTTCTTTGTTGAATCCC |
| GAPDH fwd | GCTCATCTCCTGGTACGACA |
| GAPDH rev | GTCCTTGCTCTGCATGTACT |

**Table S2: Cre/AaHIT transgenic line crosses with survival numbers.** *lox*P-AaHIT transgenic mosquitoes were crossed to tPUb-Cre mosquitoes, each being heterozygotes. The expected progeny ratios are therefore 1:1:1:1 for both transgenes, *lox*P-AaHIT only, tPUb-Cre only, and wild type. The observed numbers in each category are shown. Note the absence of any individuals carrying both transgenes. This is significant deviation from the expected outcome if all progeny classes are viable ($\chi 2$=17.47, d.f. = 3, p = 0.0006).

| group | number of larvae | % total |
|---|---|---|
| AaHIT | 15 | 30 |
| Cre | 18 | 35 |
| Cre/AaHIT | 0 | 0 |
| Wild type | 18 | 35 |

**Figure S1:** Schematic respresentation of designed plasmids

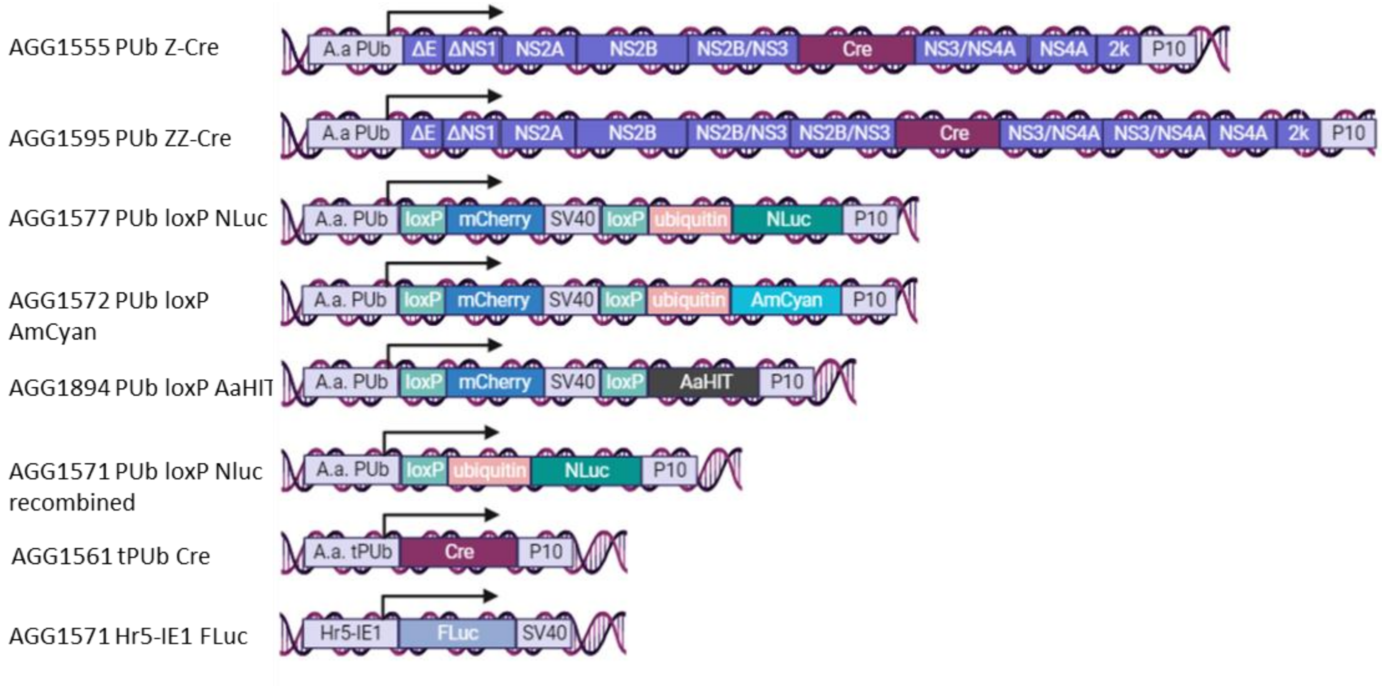
